# Structure of the Cpf1 endonuclease R-Loop complex after target DNA cleavage

**DOI:** 10.1101/122648

**Authors:** Stefano Stella, Pablo Alcón, Guillermo Montoya

## Abstract

Cpf1 is a single RNA-guided endonuclease of class 2 type V CRISPR-Cas system, emerging as a powerful genome editing tool ^1,2^. To provide insight into its DNA targeting mechanism, we have determined the crystal structure of *Francisella novicida* Cpf1 (FnCpf1) in complex with the triple strand R-loop formed after target DNA cleavage. The structure reveals a unique machinery for target DNA unwinding to form a crRNA-DNA hybrid and a displaced DNA strand inside FnCpf1. The protospacer adjacent motif (PAM) is recognised by the PAM interacting (PI) domain. In this domain, the conserved K667, K671 and K677 are arranged in a dentate manner in a loop-lysine helix-loop motif (LKL). The helix is inserted at a 45° angle to the dsDNA longitudinal axis. Unzipping of the dsDNA in a cleft arranged by acidic and hydrophobic residues facilitates the hybridization of the target DNA strand with crRNA. K667 initiates unwinding by pushing away the guanine after the PAM sequence of the dsDNA. The PAM ssDNA is funnelled towards the nuclease site, which is located 70 Å away, through a hydrophobic protein cavity with basic patches that interact with the phosphate backbone. In this catalytically active conformation the PI and the helix-loop-helix (HLH) motif in the REC1 domain adopt a “rail shape” and “flap-on” conformations, channelling the PAM strand into the cavity. A steric barrier between the RuvC-II and REC1 domains forms a “septum” that separates the displaced PAM strand and the crRNA-DNA hybrid, avoiding re-annealing of the DNA. Mutations in key residues reveal a novel mechanism to determine the DNA product length, thereby linking the PAM and DNAase sites. Our study reveals a singular working model of RNA-guided DNA cleavage by Cpf1, opening up new avenues for engineering this genome modification system^2-4^.

---

Adaptive immunity against invading genetic elements is accomplished in archaea and bacteria using ribonucleoprotein complexes composed of a combination of CRISPR associated proteins (Cas) ^5-7^ and CRISPR RNAs (crRNAs) ^8^. A Cas complex excises an invading nucleic acid into fragments, known as a protospacers, inserting them into the CRISPR repeats to form the spacer arrays. The expression and processing of these crRNAs lead to the formation of functional crRNAs that are assembled with Cas proteins to achieve interference. The resulting complexes are then guided by the crRNA to recognise and cleave complementary DNA (or RNA) sequences ^9-11^. The ability to redesign the guide RNA sequence, to target specific DNA sites, has resulted in a powerful method for genome modification in multiple biomedical and biotechnological applications. The CRISPR-Cas systems are divided into two classes and five types according to the configuration of their protein modules ^8^. The best-characterized CRISPR system is the RNA-guided endonuclease Cas9 that is categorized as class 2 type II ^12-14^. A second class 2 CRISPR system assigned to type V has more recently been identified in several bacterial genomes ^1,2,15^ extending the uses of the CRISPR family. This type V CRISPR-Cas system contains a large protein (around 1300 amino acids) termed Cpf1 (CRISPR from *Prevotella* and *Francisella;* Fig 1a, Extended Data Figure 1). In contrast with its Cas9 cousin, Cpf1 CRISPR arrays are processed into mature crRNAs without the requirement of an additional *trans*-activating crRNA (tracrRNA)^16^. Cpf1 contains an RNAase active site, which is used to process its own pre-crRNA to assemble an active ribonucleoprotein to achieve interference ^17^. The Cpf1 crRNA complex efficiently cleaves a target dsDNA containing a short T-rich PAM motif on the 5’ end of the non-target strand, in contrast to the Cas9 systems where a G-rich PAM is located on the 3’ end of the non-target strand. Moreover, Cpf1 introduces a staggered DNA double-strand break (DSB) ^2,17^ instead of the blunt end cut generated by Cas9 nucleases ^12-14^ (Fig. 1b-c), providing new capabilities for genome modification. Recently, Cpf1 enzymes have been shown to mediate robust genome editing in mammalian cells with high specificity ^3,4,18^. Therefore, this RNA-protein complex has the potential to substantially expand our ability to manipulate eukaryotic genomes.

**Figure 1.**
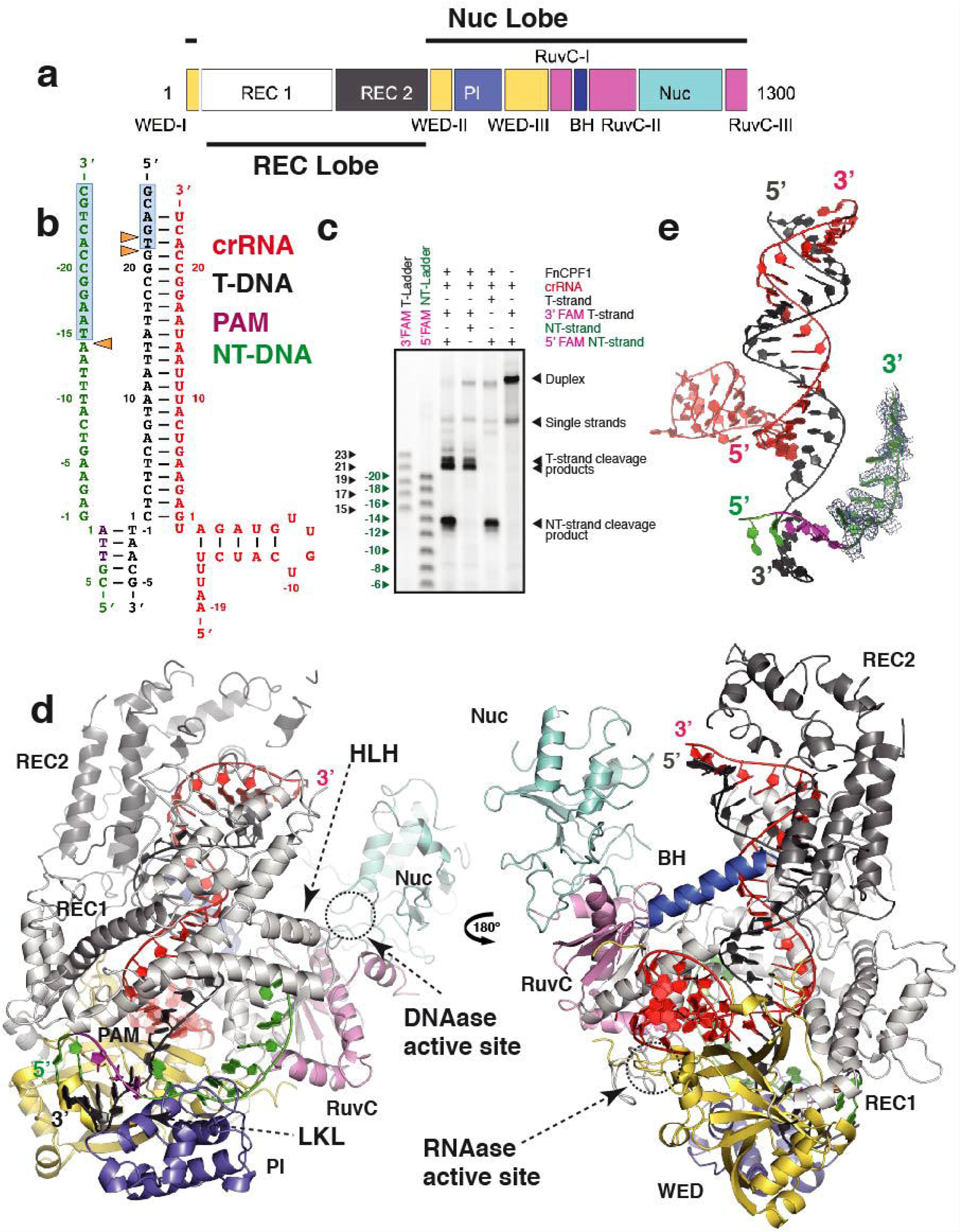
Crystal structure of the FnCpf1 R-loop complex. **a)** Domain organization of FnCpf1 protein. The polypeptide is composed by the WED (I, II and II subdomains), REC1, REC2, PI, BH, RuvC (I, II and III subdomains) and Nuc domains ^20^. The colors assigned to FnCpf1 domains are the same throughout the manuscript. **b)** Schematic diagram of the R-loop formed by the crRNA and the target DNA after cleavage of the 31-nt dsDNA target used to crystallize the complex. The colors of the DNA strands and the RNA are the same throughout the manuscript. The orange triangles = phosphodiester cleavage positions accomplished by FnCpf1 in the T and NT strands (see Fig. 1c); boxed light blue regions = section of the cleaved target DNA not present in the crystal structure. **c)** Urea TBE-PAGE gel depicting the cleavage products of the target DNA by FnCpf1. Two different ladders have been used to determine the length of the cleaved products from the target (T-ladder, black) and non-target (NT-ladder, green) DNA strands. The markers indicate the position of the cleaved phosphodiester following Fig. 1b numeration. The total number of nucleotides in the products is the marker plus the 5 extra bases of the PAM sequence (The same ladders and nomenclature are used in all the assays). The length markers show a cut at position -14 (19-nt length) of the NT-strand, while the T-strand is cleaved at position 21 (26-nt length) or 22 (27-nt length) generating an overhang in the target DNA of 7-8 nucleotides. **d)** Overview of the FnCpf1-RNA-target DNA ternary complex from the front and the back. **e)** View of the FnCpf1 R-loop structure (FnCpf1 polypeptide omitted for clarity). An omit map for the nucleotides of the NT-strand after the PAM sequence is shown at 1σ level. Unambiguous electron density is observed for 7 nucleotides of the NT-strand upstream of the PAM showing (Extended Data Figure 3).

To explore the full potential of Cpf1 for genome editing, it is vital to understand the molecular mechanisms determining its DNA targeting specificity to generate a precise DSB. The structure of *Lachnospiraceae bacterium* Cpf1 ^19^ (in complex with only 20-nt corresponding to the crRNA handle) and *Acidaminococcus sp*. Cpf1 ^20^ (including its crRNA and a target DNA with a 4-nt PAM sequence) have been solved. But these important steps toward deciphering the molecular basis of Cpf1 function provide only a catalytically inactive snapshot. Hence, questions regarding information on target DNA recognition, unzipping and the subsequent cleavage remained unanswered. Here, we present the crystal structure of a catalytically active FnCpf1 R-loop complex (Fig. 1d-e), revealing the critical residues and conformational changes in the protein to unzip and cleave the target DNA.

In order to explore the mechanism of the target DNA processing by Cpf1, we first reconstituted FnCpf1 with its crRNA *in vitro* (see Methods) to obtain a functional FnCpf1 complex. The ribonucleoprotein complex was assembled with a target dsDNA of 31 nucleotides and then purified for crystallization (Fig. 1b, see Methods). An *in vitro* analysis of the reconstituted R-loop, formed when the crRNA associated with the target DNA, revealed that the DNA overhang produced by FnCpf1 is 7-8 nucleotides (Fig. 1c), which is 2-3 nucleotides wider than previously reported ^2,17^. By increasing the concentrations of the substrate target DNA while keeping the amount of endonuclease constant, we observed a saturation of the cleavage activity, suggesting that FnCpf1 forms a stable complex with the cleaved product (Extended Data Figure 2). The crystal structure of the FnCpf1 R-loop complex was solved combining a selenomethionine derivative with a molecular replacement solution (Extended Data Table I). FnCpf1 presents an oval sea conch structure whose bilobal cavity that is formed by the Nuc and the REC lobes (Fig. 1a, d; Extended Data Video 1). The structure shows that the target DNA is cleaved yielding a triple strand R-loop (Fig. 1e, Extended Data Figure 3) with the target (complementary) DNA strand (T-strand) hybridized on the crRNA, while the dissociated PAM non-target (not complementary) DNA strand (NT-strand) is stabilized and funnelled towards the DNA nuclease site (Fig. 2a).

**Figure 2.**
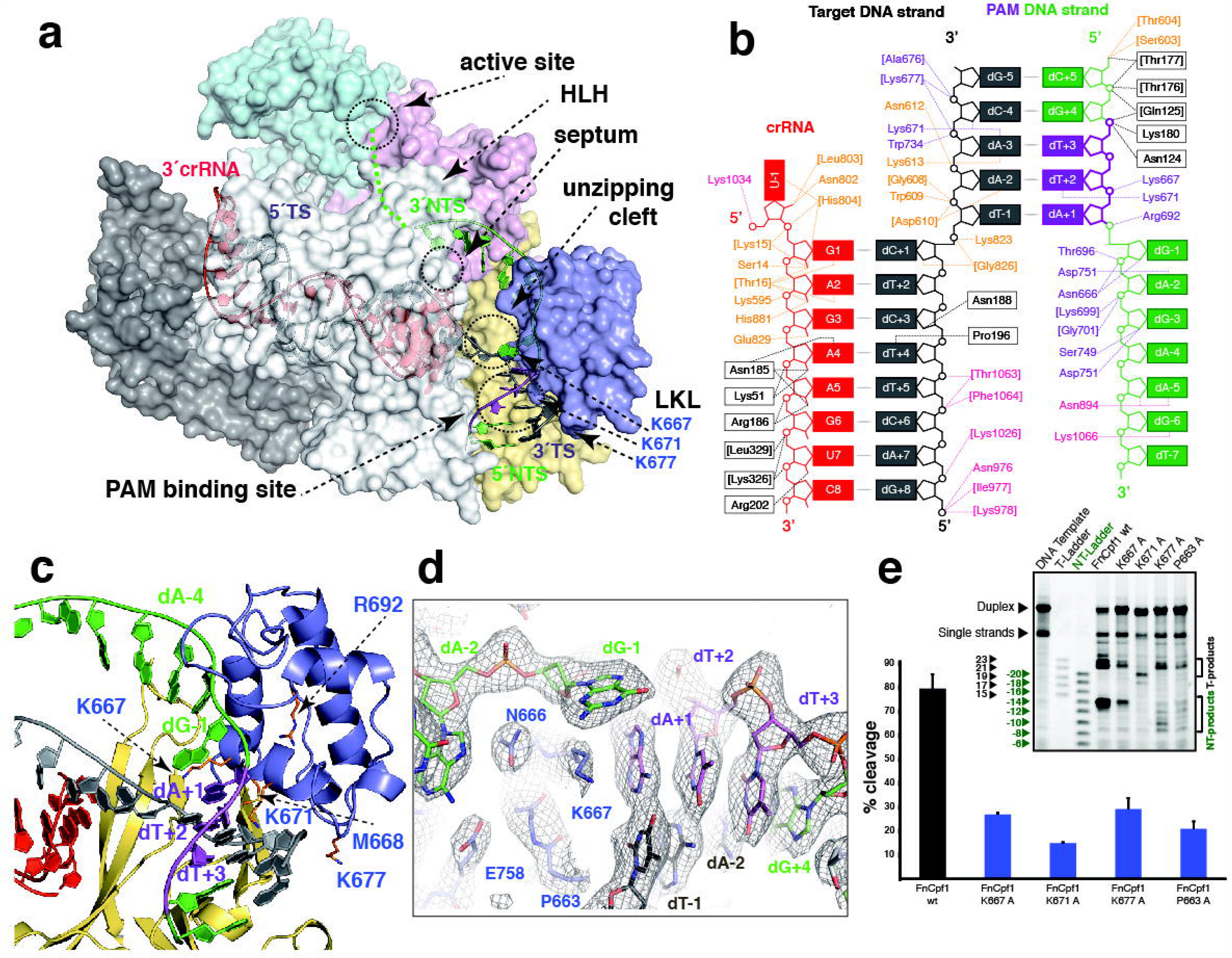
PAM recognition and uncoupling of the Watson–Crick dG-1:dC+1 pair. **a)** Surface representation of the FnCpf1 R-loop complex. The transparent surface permits visualization of the cartoon of the triple helix (crRNA-DNA hybrid and the NT-strand) inside the FnCpf1 structure. The different regions commented throughout the manuscript are indicated. The green dots show the predicted path of the NT-strand to the DNAase site after dT-7. **b)** Schematic showing key FnCpf1 R-loop interactions. For clarity, part of the crRNA-DNA hybrid has been omitted. Polar contacts of the nucleic acids with the protein side and main (in brackets) chain are indicated. The residues are labelled according to the domain colour code in Fig. 1a. **c)** Detailed view of the PAM recognition and the dsDNA separation. The conserved K667, M668, K671 and K677 (side chains have been coloured in orange for clarity) in the LKL region promote DNA unwinding. **d)** Detailed view of the dG-1:dC+1 pair uncoupling. The 2mfo-Dfc electron density refined map is contoured at 1.2σ level. **e)** Urea TBE-PAGE gel showing the cleavage activity and cleavage pattern of the P663A, K667A, K671A and K677A mutants compared with wild type (wt) FnCpf1. Each experiment has been repeated 4 times. Error bars = s.d.

**Figure 3.**
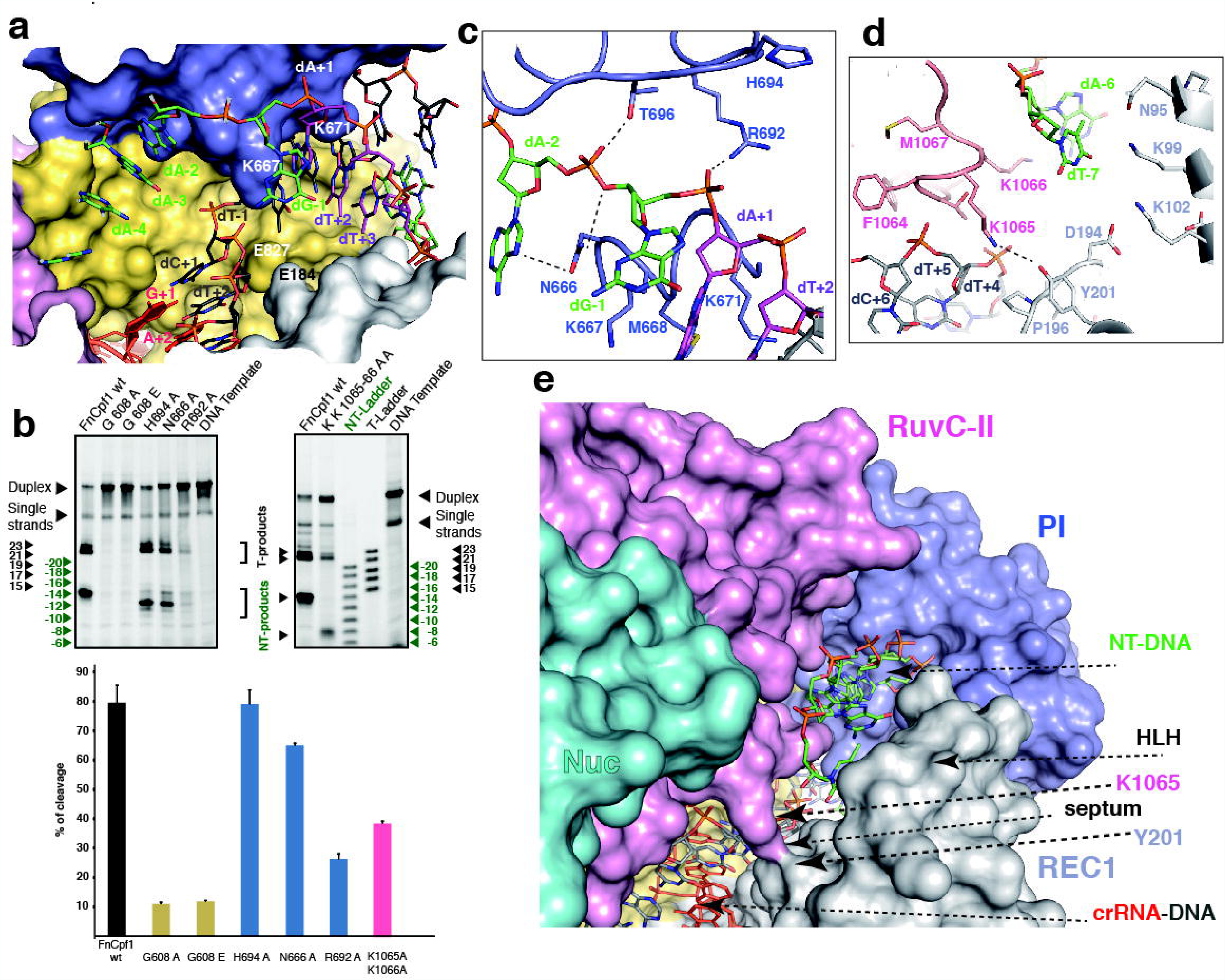
Target DNA unzipping, the PI domain “rail” conformation and the septum formation. **a)** Surface representation of the unzipping cleft formed by the WED, REC1 and RuvC domains, see also Extended Data Figure 6a. **b)** Urea TBE-PAGE gels showing the activity and cleavage pattern of mutations in the unzipping cleft, the PI domain and the septum (G608A, G608E, H694A, N666A, R692A and K1065A/K1066A) compared with wt. Each experiment repeated 4 times. Error bars = **c)** The rail conformation adopted by the PI domain in the FnCpf1 R-loop structure to accommodate and funnel the PAM NT-strand towards the active site (Extended Data Figure 7), here showing the NT-strand interaction with R692, N666 and T696. Dashed lines show the polar contacts **d)** Magnified view of the septum displaying the polar contact between the K1065 and Y201, separating the paths toward the DNAase site for the crRNA-DNA and the NT-strand. **e)** Detailed view of the “septum” observed in the FnCpf1 R-loop complex structure. The displaced NT-strand is separated from the DNA-crRNA hybrid by contacts between the REC1 and RuvC-II subdomains. The cavity is observed from the active DNAase site toward the PAM site

Unambiguous electron density was observed for seven nucleotides of the NT-strand upstream of the PAM sequence (Figure 1e and Extended Data Figure 3a). The lack of density for the rest of the nucleotides in the NT-strand indicates the high mobility on the distal end of the PAM strand as it has been shown for Cas9 ^21^. The RNA in FnCpf1 forms the handle conformation that has been observed both in the AsCpf1 ^20^ and LbCpf1 ^19^ structures. The RNA nuclease site in the WED domain (WED-III) is located at the back of the protein (Fig. 1d). The key residues, H843, K852, K869 and F873, involved in RNA processing are located in a flexible region embracing the 5´-end of the RNA handle (Extended Data Figure 4a). The alanine mutants of these residues abolished RNAase activity ^17^.

**Figure 4.**
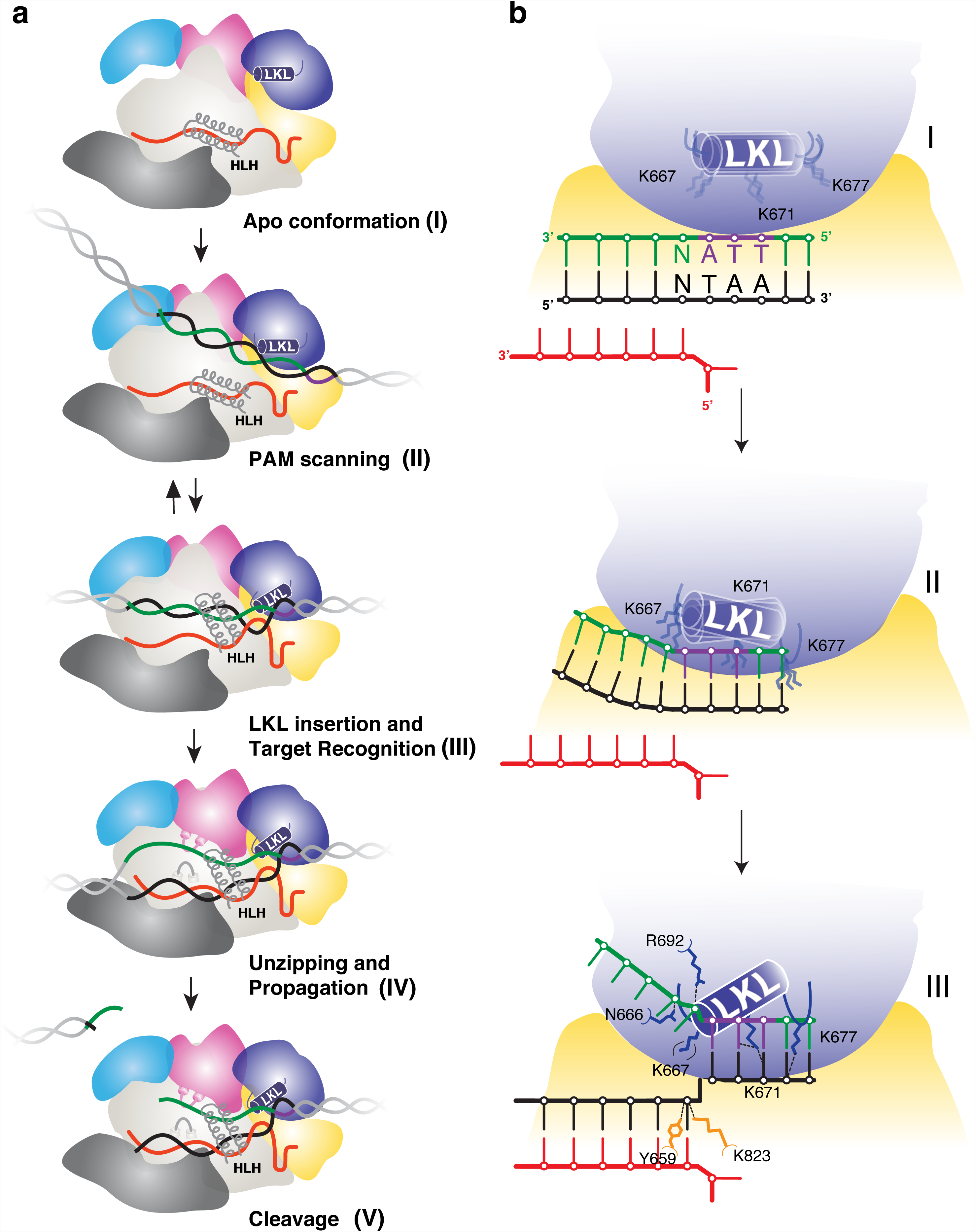
Model of PAM-dependent target DNA recognition and ATP-independent unwinding by Cpf1. **a)** The ribonucleoparticle is in a relaxed open conformation (I), until the presence of possible target DNA substrates promotes a compact conformation that allows the scanning of PAM sequences (II). Binding to a PAM sequence promotes the insertion of the helix in the LKL region and the “flap-on” conformation of the HLH motif (III). If coupling between the nucleotides of the T-strand and the crRNA is completed in the unzipping cleft, the PI domain adopts the “rail” conformation while the NT-strand is displaced and the “septum” between the REC and Nuc lobes is formed (IV). At the active site, catalysis proceeds generating a DNA overhang of 7-8nt (V). **b)** Detailed view of the LKL region during PAM scanning, recognition and NT-strand displacement, to show the role of the conserved K667, K671 and K677 residues. The LKL region scans the DNA looking for a PAM sequence (I). Once a candidate is found K677 contacts the phosphate before the first base in the PAM site in the T-strand allowing K671 to test binding in the central part of the PAM sequence contacting the bases in positions dA-3 and dT+2 (II). In the case that K671 is properly positioned, The PI domain adopts the rail conformation and the helix in the LKL region is inserted in the dsDNA. The Watson–Crick base pair after the PAM is uncoupled by K667 and the inversion of the phosphate in the -1 nucleotide by K823 and Y659 in the T-strand, thus starting the unzipping of the dsDNA to allow hybridization with the crRNA. The residues R692 and N666 stabilize the displaced NT-strand by contacting the -1 and -2 phosphates of the backbone (III).

PAM sequence recognition is a critical aspect of DNA targeting by RNA guided nucleases targeting, as it is a prerequisite for ATP-independent DNA strand separation and crRNA–target-DNA heteroduplex formation ^22^. In contrast with Cas9, the PAM binding site in FnCpf1 is IZ70 Å away from the nuclease active site, resulting in the physical separation of the target recognition and cleavage sites in the ribonucleoparticle (Fig. 2a). FnCpf1 recognises a three nucleotide (5′-TTN-3′) PAM sequence, while AsCpf1 and LbCpf1 require a four nucleotide (5′-TTTN-3′) PAM ^2^. The WED (WED-II and III), REC1 and PI domains build the binding region of the PAM sequence (Fig. 2a-b). The side and main chains of residues belonging to the WED-II, WED-III and REC1 domains make many polar and van der Waals contacts (Fig. 2b), while a loop-lysine helix-loop motif (LKL motif comprising residues L662 to I679) in the PI domain contains the conserved lysine residues, K667, K671 and K677 (Extended Data Figure 1).

We found that the LKL region has an essential role in recognizing PAM and promoting strand separation (Fig. 2c). Only K671 interacts with the bases in the PAM region, as is the case for AsCpf1^20^, suggesting that shape might play a more important role than direct base readout in PAM recognition. Our structure shows that the helix in the LKL region plays a key role in dsDNA unzipping to promote hybridization between the T-strand and the crRNA, by displacing the PAM NT-strand (Fig. 2c). The helix is inserted at an angle of 45° with respect to the dsDNA longitudinal axis, working as a dentate plough unwinding the helical dsDNA (Extended Data Figure 4b). The LKL element inserts the three lysine “pins” (K667, K671 and K677) in a staggered manner on the side facing the DNA (Fig. 2c, Extended Data Figure 4b). K677 is inserted in the major groove of the DNA interacting with the backbone of 3´- dG-5 in the T-strand, while K671 is inserted in the minor groove contacting the oxygen of the C2 carbonyl in dT+2, the N3 of dA-3 in the PAM sequence and the oxygen of the deoxyribose of dC-4 (Fig. 2b). Finally, we observed that the side chain of K667 disrupts the Watson–Crick interaction of the dG-1:dC+1 pair after the PAM sequence, pushing the dG-1 away from the dsDNA conformation (Fig. 2d). The uncoupling of the first base pair of the target DNA initiates the unzipping of the double strand facilitating the hybridization of the T-strand with the crRNA.

Other conserved residues in the LKL region such as P663 and P670, collaborate to insert the LKL plough and unzip the target DNA. P663 induces a conformation in the loop that positions the helix in the correct orientation for its insertion in the dsDNA; accordingly, the P663A mutation severely reduces FnCpf1 cleavage activity (Fig. 2e). The other conserved proline (P670) in the helical section of the LKL region, favours the insertion of K671 in the PAM sequence, and the conserved M668 also contributes by disrupting the base pairing of the double helix, thus facilitating the uncoupling of dG-1 by the invariant K667 (Fig. 2d). As observed in AsCpf1^20^, the K671A mutation in FnCpf1 presents a large activity reduction in agreement with its important role in PAM recognition. Remarkably, we found that the other lysine mutants, K667A and K677A, also reduce FnCpf1 DNA cleavage activity around 70%, indicating the pivotal role of the three lysines in PAM recognition and unzipping of the target DNA (Fig. 2e).

In addition to the activity decrease, we observed that while the K667A mutant displays DNA cleavage products similar to the wild type, K677A generates a different cleavage pattern, in which the NT-strand is cleaved into shorter fragments than that produced by the wild type protein (Fig. 1c, Fig. 2e, Extended Data Figure 5). The same pattern can be observed for the P663A mutant (Fig. 1c, Fig. 2e, Extended Data Figure 5). This suggests that P663 and K677 initiate the positioning of the LKL helix, thereby facilitating the insertion of K671 and K667 in the target DNA and thus determining the length of the cleaved product of the NT-strand. Our structure– function analysis suggests that positioning of the LKL region may function as the marker of an internal molecular ruler linking the 70 Å distant PAM and DNAase active sites to hydrolyse the correct phosphodiester bond.

Overlapping with the region where the first base pair of the target DNA is uncoupled, the WED (WED-I-II-III), REC1 and RuvC (RuvC-I) domains build the unzipping cleft (Fig. 2a, 3a). This cleft is composed of the well conserved E184, P883, I884, G826, A828, E827, E829, F831, H881, G608, L661, L662 and P663 (Extended Data Figure 1 and Extended Data Figure 6a). The FnCpf1 R-loop structure suggests that the electrostatic properties of the arrangement of the acidic and hydrophobic residues in the unzipping cleft facilitate the coupling of the dC+1 of the T-strand of the DNA with the G+1 of the crRNA (Fig. 3a, Extended Data Figure 6a). The conserved E184 and E827, which are positioned behind the T-strand, facilitate the scanning of the complementary base and the formation of the crRNA-DNA hybrid by electrostatic repulsion of the phosphate backbone of dC+1 and dT+2 in the T-strand (Extended Data Figure 6a). The phosphate of dT-1 of the T-strand is inverted by the interaction with the side chains of K823, Y659 and the main chains of G826 and N825, which also house the dT-1 phosphate. Mutation of the conserved G826 in AsCpf1 severely disturbs phosphate inversion ^20^. We found that mutations to alanine and glutamate in the neighbouring G608 residue also severely affect activity (Fig. 3b, Extended Data Figure 5). This residue contacts the PAM dA-2 phosphate, and is located in the cleft close to the PAM binding site. The modelling of any other side chains in those positions results in steric hindrance, which indicates that the DNA could not be accommodated. Thereby, these two invariant glycine residues are essential for housing the target DNA.

A comparison of the LbCpf1 apo ^19^ and FnCpf1 R-loop structures indicates that Cpf1 undergoes a large conformational rearrangement in the PI domain (Extended Data Figure 7) and the REC lobe (Extended Data Figure 8). We observe that the PI domain in the FnCpf1 R-loop complex undergoes a readjustment to accommodate the NT-strand. The nucleotides of the NT-strand exhibit an extended distorted helical conformation, and the phosphate backbone displays a number of contacts with residues of the PI and RuvC (RuvC-I and II) domains (Fig. 2b). The residues contained in the loop of the PI domain comprising residues P670 to E715 adopt a “rail shape”, embracing the NT-strand (Fig. 3c, Extended Data Figure 7), and guiding it into a hydrophobic cavity with basic patches to the active site (Extended Data Figure 6b). The R692 residue contacts the dG-1 phosphate from the topside of the phosphate backbone of the NT-strand, while N666, which is inserted between dA-2 and dG-1, and interacts with the dA-2 phosphate from the underneath (Fig. 3c). T696 also interacts with the phosphate backbone of the NT-strand, funnelling the ssDNA to the active site located between the Nuc and RuvC domains of FnCpf1. A R692A mutation greatly reduces the endonuclease activity of FnCpf1 by 70%, suggesting that this first residue in the “rail” has a key role interacting with the uncoupled dG-1 and directing the NT-strand to the DNAase site. Although the N666A and H694A mutants produce only a small reduction in activity (Fig. 3b, Extended Data Figure 5) (5-15%), their cleaved NT-strand products are two bases shorter than the wild type (Fig. 3b), indicating that in addition to the LKL region, the residues in this loop of the PI domain are also involved in the internal molecular ruler determining the length of the NT-strand to cleave the correct phosphodiester bond.

Some of the conformational differences observed between the apo^19^ and the R-loop complex structures in the distal REC2 subdomain could be attributed to the lack of a 23-nt segment of the guide RNA (Extended Data Figure 8a). However, a superposition of the structures suggests that changes in the HLH region of REC1 (F61-F123) could be induced by the binding of the PAM sequence to protect the NT-strand after unzipping (Extended Data Figure 8b and Extended Data Video 2). The HLH segment in the apo conformation would be in contact with the well conserved T197-S220 and F264-G277 helices in the REC1 subdomain like in the LbCpf1 apo conformation (Extended Data Figure 1 and 8b). The HLH segment is in a relaxed conformation aligned with the REC lobe in the apo structure (Extended Data Figure 8b). The binding of the target DNA induces a closure of the protein, which positions the Nuc and REC lobes closer ^23^. The W971 in the BH domain would be anchored in the REC2 subdomain hydrophobic pocket, as observed in AsCpf1^20^, thus stopping the movement and locking the closed active conformation. Hence, the FnCpf1 R-loop structure suggests that PAM recognition and NT-strand displacement might induce the HLH segment to abandon its resting position, rotating 45° as a rigid segment mimicking a “flap-on” movement (Extended Data Video 2). The HLH segment “flap-on” and the PI “rail” conformations would protect the NT-strand by partially occluding the cleft where it is located, directing it to the DNAase site for phosphodiester cleavage (Extended Data Figure 7 and 8).

While examining the FnCpf1-crRNA-target DNA ternary complex in more detail, we observed that there were two separate paths in its bilobal architecture starting from the PAM binding site. The crRNA-DNA hybrid spans all the length of the central protein cavity between the REC and Nuc lobes, while the displaced NT-strand runs along a second trail separated from the previous one by contacts between the REC1 (T197-V204) and RuvC (F1061-Q1070) domains. These conserved segments interact forming a thin separation, the “septum”, which establishes a steric barrier at the level of the dT-7 of the NT-strand, thus avoiding possible re-annealing with the complementary DNA strand (Fig. 3d-e). K1065 and Y201 make a polar interaction between the NH_2_ and the OH of their side chains physically separating the NT-strand and the crRNA-DNA hybrid. This second path is observed only in the presence of the displaced NT-strand (Fig. 3e). The mutation of the two consecutive lysines (K1065A/K1066A) in the RuvC-II subdomain showed a 52% activity decrease, revealing the importance of the septum in Cpf1 (Fig. 3b, Extended Data Figure 5). Remarkably, the cleavage pattern of the NT-strand displays a 6-nt shorter product, while the T-strand is cut at the same position as the wild type, indicating that the different regions building the path involved in directing the NT-strand to the DNAase site are essential for correct cleavage (Fig. 3b).

Understanding the mechanism by which Cpf1 recognizes, unzips and cleaves target DNA is vital to fully exploiting the potential of this system for genome editing. Here, we have shown how FnCpf1 catalyzes the R-loop structure through structural changes leading to precise DNA cleavage and generation of a DSB. Our findings suggest a working model (Fig. 4a) by which the apo protein-RNA complex first undergoes a conformational change from a relaxed open conformation (I) to a compact DNA bound conformation (II) in order to scan target DNAs looking for complementary sequences to its crRNA. The recognition of the PAM sequence promotes the “flap-on” conformation of the HLH motif (III). After PAM recognition, the helix in the LKL region is inserted in the target dsDNA, starting the melting of the dsDNA and the uncoupling of the bases proximal to the PAM. K671 and K677 interact with the PAM sequence, and K667 promotes the uncoupling of the first Watson–Crick base pair after the PAM sequence (Fig. 4b), the phosphate in the nucleotide in -1 position of the T-strand is inverted and the dsDNA is unwound in the unzipping cleft, thus allowing the pairing of the initial uncoupled bases of the T-strand with the crRNA (III). The electrostatic environment in the cleft favours the pairing of the initially uncoupled DNA nucleotides with the crRNA. If a mismatch occurs and the energetics of the process do not favour the crRNA-DNA hybrid coupling, the helix in the LKL region is removed, the two strands hybridize, and Cpf1 continues searching for a PAM sequence containing the complementary target DNA (II), as has been proposed for other RNA-guided endonucleases ^24,25^. In case of complementarity, once the initial Watson–Crick coupling between the bases after the PAM is completed in the unzipping cleft, the PI domain adopts the “rail” conformation while the NT-strand is displaced and the “septum” between the REC and Nuc lobes is formed (IV). Consequently, propagation will result in the crRNA-DNA hybrid residing in the main cavity of Cpf1 (IV), leading to the generation of a DSB in the target DNA (V). Our data suggest that FnCpf1 remains associated to the DSB after cleavage and that there is a 7-8-nt 5’ overhang produced in the target DNA (Fig. 1c and Extended Data Figure 2).

Our analysis suggests that the changes in the PI domain (rail conformation) and HLH segment (flap-on conformation) in REC1, together with the more compact structure of Cpf1, generate a “septum” providing a second path inside the protein that is used to direct the NT-strand toward the DNAase active site. Therefore, in conjunction with the other structural features, the PI domain and, particularly the LKL region, could act as molecular ruler determining the appropriate length to the distant DNAase site for correct cleavage of the target DNA, thus linking the molecular events of recognition and catalysis.

Our findings provide a detailed understanding into Cpf1 endonucleases, which could have important implications in genome modifying applications. Cpf1 can be harnessed to gene targeting in mammals^3,4,26^, and may be also adapted for other possible applications such as “base editing”^27^ or large-scale genetic studies ^28^. Thus our results provide valuable insight into Cpf1 RNA-guided DNA cleavage, establishing a detailed outline for rational engineering, thus widening the CRISPR repertoire.

## Acknowledgements

The Novo Nordisk Foundation Center for Protein Research is supported financially by the Novo Nordisk Foundation (Grant NNF14CC0001). We thank the beamline staff of X06SA at the Swiss Light Source (Paul Scherrer Institut, Villigen, Switzerland), the EMBL beamline P14 at DESY (Hamburg, Germany) and the MAX-Lab in Lund for support during data collection. Data processing and the calculation of the structures have been performed in the Computerome (Danish National Computer for Life Sciences). We also thank Havva Koc and Motiejus Melynis from the Prokaryotic Team of the Protein Production Facility Platform at CPR for the excellent technical assistance.

### Author contributions

SS expressed and purified the FnCpf1 R-loop complex with the help of PA. Both prepared the crystals and performed the biochemical experiments. All authors collected diffraction data. GM performed data processing and crystallographic analysis. All authors were involved in model building and refinement. All authors discussed the data, and helped GM to write the manuscript and prepare the figures. The coordinates and structure factors for the FnCpf1 R-loop complex have been deposited in the PDB under accession number XXX (waiting for the PDB code).

### Competing Financial Interests

The authors declare no competing financial interests.

## Methods

### Protein expression and purification

The gene encoding for Cpf1 from *Francisella novicida* U112 (FnCpf1) was obtained from Addgene (Item ID: 69975). The ORF was amplified by PCR using the primers FnCpf1-Forward and FnCpf1-Reverse (Extended Data Table II) and cloned into pET21a. The obtained construct was transformed in *Escherichia coli* BL21 star (DE3) cells containing the pRare2 plasmid. Protein expression was induced with 1 mM IPTG for 3 hours. For the preparation of selenomethionine-substituted protein, the cells were grown in SelenoMethionine Medium Complete (Molecular Dimensions) including 40 μg ml^−1^ selenomethionine and the expression was induced as described above. Cells were resuspended in lysis buffer (50 mM Bicine pH 8.0, 150 mM KCl, 1 tablet/50 ml Complete Inhibitor cocktail EDTA Free (Roche), 50 U/ml Benzonase, 1 mg/ml lysozyme and 0.5 mM TCEP). After cell disruption by French press, cell debris and insoluble particles were removed by centrifugation at 10,000g at 4°C. The supernatant was loaded onto a HisTrap column (GE Healthcare) equilibrated in buffer A (50 mM Bicine pH 8.0, 150 mM KCl and 0.5 mM TCEP). After the sample was loaded, the column was washed with buffer A containing 5 mM imidazole to prevent non-specific binding of contaminants to the resin. The sample was eluted with 50 mM Bicine pH 8.0, 150 mM KCl, 0.5 mM TCEP and 250 mM imidazole. Enriched protein fractions were pooled together and applied onto a HiTrap Heparin HP column (GE Healthcare) equilibrated with buffer A. The protein was eluted with a linear gradient of 0-100% buffer H (50 mM Bicine pH 8.0, 1 M KCl and 0.5 mM TCEP). Protein-rich fractions were collected and concentrated (using 100 kDa MWCO Centriprep Amicon Ultra devices) and subsequently loaded onto a HiLoad 16/60 200 Superdex column (GE Healthcare) equilibrated in buffer A. The protein peaks were concentrated (using 100 kDa MWCO Centriprep Amicon Ultra devices), flash-frozen in liquid nitrogen and stored at -80°C. The protein concentration was determined using the theoretical molar extinction coefficient at 280 nm calculated from the amino-acid composition. An overloaded SDS–PAGE gel stained with SimplyBlue (Invitrogen) displayed a highly pure protein preparation.

### RNA production and purification

DNA oligonucleotides corresponding to the reverse complemented sequence of the target site (RNA template) and a short T7 priming sequence (T7 primer) were purchased from Integrated DNA Technologies (IDT; Extended Data Table II). The oligonucleotides were annealed at a final concentration of 20 μM in annealing buffer containing 150 mM KCl by heating the mixture up to 95°C for 10 min followed by a cool ramp to 4°C over 10 min. This partial DNA duplex was used as template in the transcription reaction carried out with HiScribe T7 Quick High Yield RNA Synthesis (NEB). The reaction was stopped using 2X stop solution (50 mM EDTA, 20 mM Tris-HCl pH 8.0 and 8 M Urea) and the RNA was denatured at 95°C for 10 min. The transcription product was purified by preparative electrophoresis with a Bio-Rad Model 491 PrepCell apparatus equipped with the 37mm i.d. gel tube using a 9cm tall 1X TBE, 15% (19:1) polyacrylamide/7 M urea gel at room temperature.

### FnCpf1 R-loop complex formation

The 31-nucleotide DNA template was annealed using a single strand oligonucleotide, according to the protocol above for the preparation of the transcription template. The complex was assembled in reconstitution buffer consisting of 150 mM KCl, 50 mM Bicine (pH8.0), 5 mM MgCl_2_ at a molar ratio of protein:RNA:DNA 1:1.3:1.7. The FnCpf1 protein was mixed with crRNA and incubated for 30 min at 25°C before adding the 31bp target DNA duplex (NT-strand and T-strand Extended Data Table II) followed by one-hour incubation at 25°C. The reconstituted FnCpf1 R-loop complex was further purified by preparative electrophoresis with a Bio-Rad Model 491 PrepCell apparatus using a 5% (w/v) non-denaturing polyacrylamide gel at 4°C. The highly pure and homogeneous complex was separated from free DNA and high-molecular weight aggregates and immediately concentrated to 7 mg/ml using a Vivaspin(r) 20 50000 MWCO.

### Crystallization

Initial crystallisation screening was performed at 20°C using the sitting-drop vapour-diffusion method and testing a collection of commercially available crystallisation screens. After five days of incubation, the extensive initial screening rendered plate-like crystals in 0.35 M sodium thiocyanate, 20% w/v PEG 3350. Following the initial hit identification, crystal growth was optimised using a Dragonfly screen optimiser. Selenomethionine-modified crystals were obtained under the same conditions.

### Structure determination, model building and refinement

The structure of FnCpf1 R-loop complex was determined by combining a molecular replacement solution, using AsCpf1^20^ as a search model in a native data set (λ=1.00), and a FnCpf1 selenium derivative crystal, where a single-wavelength anomalous diffraction (SAD) data set was collected at the peak of Se K absorption edge (λ=0.978). Both the native and the SAD data were collected from frozen crystals at 100 K using an EIGER detector at the PXI-XS06 beamline (Swiss Light Source Villigen, Switzerland). Data processing and scaling were accomplished with *XDS* ^29^ and *AIMLESS* ^30^ as implemented in *autoPROC* ^31^ (Extended Data Table I). All methionines were substituted by selenomethionine and 10 out of the 13 possible Se sites were identified using *SHELX* package ^32^. Initial phases were calculated at 3.7 Å resolution *PHASER* as implemented in the *PHENIX* suite ^33,34^. These initial phases were extended to 3.0 Å resolution using the native data set with the PHENIX *Autobuild* routine. The initial Cα model was remodelled manually with Coot ^35^ and O 36,37 and refined initially using PHENIX ^33^. Se anomalous maps were used as a guide during model building. Several rounds of manual building and refinement using BUSTER ^38^ yielded the refinement and data collection statistics summarized in the Extended Data Table I. The identification and analysis of the protein-DNA-RNA hydrogen bonds and van der Waals contacts were performed with the Protein Interfaces, Surfaces and Assemblies service (PISA) as implemented in the CCP4 suite 39 and Coot ^35^. The figures for the manuscript have been produced with PyMOL (The PyMOL Molecular Graphics System, Version 1.3, Schrödinger, LLC). The final model has an Rwork/Rfree of 23/26% with good stereochemistry according to MolProbity (Extended Data Table I) and only 0.7% of the residues in disallowed regions of the Ramachandran plot.

### Mutagenesis

Mutagenesis was performed using the Q5 site direct mutagenesis kit (NEB). The primers were designed according to the NEBaseChanger (see Extended Data Table II). The FnCpf1 mutants were expressed and purified as described above. The purity of mutant proteins was analysed by SDS-PAGE stained with SimplyBlue (Invitrogen; Extended Data Figure 5).

### In vitro cleavage assay

The fluorescently labelled DNA substrate was prepared by annealing the T-strand-6FAM-A and NT-strand-6FAM-A (DNA1A) under the same condition described above (Extended Data Table II). To identify which strands were cleaved the DNA substrate was prepared by annealing NT-strand-6FAM-A with T-strand-A (DNA2A) and T-strand-6FAM-A with T-strand-A (DNA3A) (Extended Data Table II). The RNA/protein complexes were formed in 20mM Bicine-HCl pH8, 150mM KCl, 0.5mM TCPE pH8 and 5 mM MgCl using 6 nmoles of purified RNA and 4 nmoles of purified protein. The complex was incubated at 25°C for 30 min before adding 2 nmoles of DNA substrate. The mixture was incubated at 25°C for 1 hour and the reaction was stopped by adding equal volume of stop solution (8 M Urea and 100 mM EDTA at pH8) followed by incubation at 95°C for 5 min. The samples were loaded on 15% Novex TBE-Urea Gels (Invitrogen) and run according to the manufacture instructions. The gel was visualized using an Odyssey FC Imaging System (Li-Cor) and the intensity (I) of the DNA bands was quantified using ImageStudio. The percentage of DNA cleavage was calculated using the following formula: I_(T-strand product)_ + I_(NT-products)_ /I _(total DNA)_. The total DNA was calculated as the sum of I_(duplex)_ + I_(single strands)_ + I_(T-strand products)_ +I _(NT-products)_.

### In vitro saturation assay

The DNA substrate was prepared by annealing the oligo T-strand-6FAM and NT-strand-6FAM (DNA1) under the same condition describe above. To identify which strand was cleaved the DNA substrate was prepared by annealing NT-strand-6FAM with T-strand (DNA2) and T-strand-6FAM with T-strand (DNA3) (Extended Data Table II and Extended Data Figure 2). The RNA/FnCpf1 complex was prepared as described above and incubated with increasing amounts of DNA 1 from 60 pmoles to 60 nmoles. The reactions were incubated at 25°C for 3 hours, and then stopped and loaded on the gels as described above.

